# Selective targeting of *Plasmodium falciparum* hexose transporter by phytochemical Ginsenoside Rg1 disrupts glucose metabolism and blocks development of parasite

**DOI:** 10.1101/2025.05.23.655744

**Authors:** Sadat Shafi, Preeti Maurya, Ruby Bansal, Jyoti Sharma, Sai Kumar Mishra, Abul Kalam Najmi, Shailja Singh

## Abstract

The emergence of resistance to first-line antimalarial therapies highlights the critical need for next-generation drugs that target distinct molecular pathways and employ novel mechanisms of action. Notably, the intra-erythrocytic parasite development is highly dependent on a sustained glucose supply as their fundamental energy source. Therefore, exploiting a “selective starvation” strategy, by targeting the parasite’s reliance on glucose metabolism, particularly through the *Plasmodium falciparum* hexose transporter (*Pf*HT1), which is critical for parasite survival can serve as a promising therapeutic approach to combat multidrug-resistant *Plasmodium* parasites. Through molecular docking and structure-based drug design approach, we identified a natural compound, Ginsenoside Rg1 (G-Rg1) from drug bank database library, as a potential *Pf*HT1 inhibitor. The *Pf*HT1 specificity of G-Rg1 was validated using yeast complementation model. Subsequently, to investigate the role of *Pf*HT1 in drug resistant *Pf* parasites we investigated the stage-specific expression of *Pf*HT1 in both artemisinin (ART)-sensitive and resistant *Pf* parasites and reported its elevated expression in resistant parasites, predicting its role in their survival. Notably, *in vitro* growth inhibition studies demonstrated that G-Rg1 effectively suppressed the growth of both ART-sensitive and resistant *Pf* parasites. Additionally, G-Rg1 potentiated the efficacy of dihydroartemisinin in combination and ring survival assays, indicating its potential to circumvent resistance mechanisms. G-Rg1 administration alone and in combination with ART, in *P. berghei* ANKA-infected mice reduced parasite multiplication and increased mean survival time. Our findings support G-Rg1 as a promising candidate for drug development against malaria, highlighting the potential of targeting *Pf*HT1 to combat drug-resistant malaria.

## INTRODUCTION

Despite extensive efforts over several decades to eliminate malaria, it is still a prevalent endemic in the tropical and subtropical subparts of Southeast Asia and sub-Saharan Africa (1). The causative agent responsible for malaria is *Plasmodium* parasite, which is disseminated through the bite of female *Anopheles* mosquito, with *Plasmodium falciparum* (*Pf*) listed as the most lethal form (2). Although various classes of drugs, including artemisinin (ART) combination therapies (ACTs), have been developed successfully for malaria treatment, nevertheless the rise of multi-drug-resistant *Plasmodium* strains has created a critical requirement to search and therapeutically investigate new targets to effectively combat this deadly disease (3).

To discover new anti-malarial targets, identifying mechanisms crucial for parasite survival is essential. Notably, glucose represents an important source of energy for *Pf* parasites (4), as they are highly dependent on glycolysis to produce adenosine triphosphate (ATP) for their energy needs, with the more efficient tricarboxylic acid cycle being mostly inactive during the asexual blood stage (5). Lacking significant intracellular energy reserves throughout most of their lifecycle, these parasites depend on a continuous supply of glucose from their hosts as their main energy source (6). Consequently, infected erythrocytes exhibit approximately a 100-fold increase in glucose acquisition compared to un-infected erythrocytes (7, 8). This metabolic dependence of parasites on glucose highlights metabolic pathways, including the glucose transporter, as an attractive therapeutic target (9). *Pf* hexose transporter (*Pf*HT) serves as the primary glucose transporter in *Pf*, expressed by a single-copy gene without closely related paralogues. *Pf*HT1 facilitates the acquisition of this vital nutrient across the plasma membrane of parasite, ensuring the parasite’s survival (10). Genetic studies have confirmed *Pf*HT1’s essential role in *Plasmodium* species, and it has been independently validated as a promising drug target for malaria treatment (11). Previously, the small-molecule glucose derivative compound 3361 (C3361) has been shown to inhibit *Pf*HT1, effectively reducing the growth of intra-erythrocytic *Pf* parasites *in vitro* (12, 13). This supports the notion of *Pf*HT1 as a viable target for anti-malarial drugs. Based on this hypothesis, employing a structure-based drug design (SBDD), approach to screen drug libraries for compounds that inhibit *Pf*HT1 may contribute to the identification of new anti-malarial therapeutics.

Phytocompounds serve as a valuable resource for current drug research. They have been used extensively as supplemental therapy to address parasite diseases (14, 15). Their source, accessibility, safety, and cost-effectiveness render them an acceptable alternative to contemporary medicine. Notably, many current anti-malarials including quinine and ART have been obtained from herbal sources . Therefore, in this study, we screened drug bank database and utilized InstaDock to find high-affinity *Pf*HT1 binding partner by molecular docking. We identified Ginsenoside Rg1 (G-Rg1) as a potential inhibitor of *Pf*HT1. Notably, scientific evidence has predicted G-Rg1 as the most effective component of *Panax ginseng* with pleiotropic therapeutic effects, including anti-inflammatory, immune-modulatory, anti-cancer, neuroprotective, and anti-microbial activity (16–18). In line with this, we proposed ‘selective metabolic starvation’ by inhibiting *Pf*HT1 as an effective therapeutic approach in combating drug resistant *Plasmodium* infection and evaluated anti-malarial potential of G-Rg1 in pre-clinical settings.

In order to identify the *Pf*HT1 specificity of G-Rg1, we used yeast complementation model system. Like plasmodium, yeast (*Saccharomyces cerevisiae*) requires constant supply of glucose to grow and yeast hexose transporter (hxt) is crucial for its glucose uptake. We used a yeast strain (Δhxt) where all the glucose transporters have been deleted, limiting its growth on glucose media. This strain provides a platform to clone and characterize hexose transporters from other species. Functional complementation of *Pf*HT1 in *S. cerevisiae* hxt mutants (Δhxt) rescued the growth of yeast significantly, but displayed diminshed growth in the presence of G-Rg1. This confirmed the *Pf*HT1 inhibitory potential of G-Rg1. Furthermore, to extend our observations to the role of *Pf*HT1 in drug resistant *Pf* parasites, we demonstrated the stage specific expression of *Pf*HT1 in ART-sensitive and resistant parasites. Quantitation of mRNA encoding this transporter revealed that its expression is elevated in mature stages of resistant parasites, thereby identifying it as a potential drug target to combat multidrug-resistant *Pf* parasites. Upon experimental validation, we found G-Rg1 exhibited potent anti-parasitic activity against *Pf* parasites in both *in vitro* assays and *in vivo* mice model. G-Rg1 suppressed the growth of ART-sensitive and resistant parasites within a sub-micromolar concentration, which was confirmed by glucose uptake blocking effect of G-Rg1 in parasites. Moreover, G-Rg1 significantly decreased the ring-stage growth and survival of *Pf* parasites resistant to ART (*Pf*Kelch13^R539T^). Subsequently, G-Rg1 treatment improved the efficacy of dihydroartemisinin (DHA) in inhibiting parasite proliferation in combination and ring survival assays (RSA), indicating that targeting the parasite’s resistance mechanism is a promising approach to combat the rapid development of drug-resistant parasites. In *in vivo* studies, administration of G-Rg1 alone and/or with ART markedly suppressed the parasite burden and elevated mean survival in infected mice, compared to untreated mice. Overall, our results provide strong evidence for the antimalarial potential of G-Rg1 and support the therapeutic relevance of targeting *Pf*HT1 in the development of novel antimalarial drugs.

## RESULTS

### Ginsenoside Rg1 as a potential binder of *P. falciparum* hexose transporter

The molecular docking simulations revealed a distinct binding mode for G-Rg1 within the *Pf*HT-1 binding pocket. Schematic illustration of complex formation between G-Rg1 (2D structure depicted in **Figure 1A**) and the binding pocket of *Pf*HT1 are shown in **Figure 1B (i)** and **(ii)** respectively. The ligand of interest, G-Rg1, showed a binding energy (ΔG_bind_) of -8.2kcal/mol in its most favorable docked pose. Analysis of the protein-ligand interactions showed that G-Rg1 primarily engaged with a pocket formed by residues Asn48, Lys51, Thr74, Ser315, Ser317, Asn318, and Glu319 **[Figure 1C (i)]**. Importantly, all of these interactions were characterized by hydrogen bonding, likely contributing to the compound’s strong binding affinity to *Pf*HT-1.

**Figure 1:**
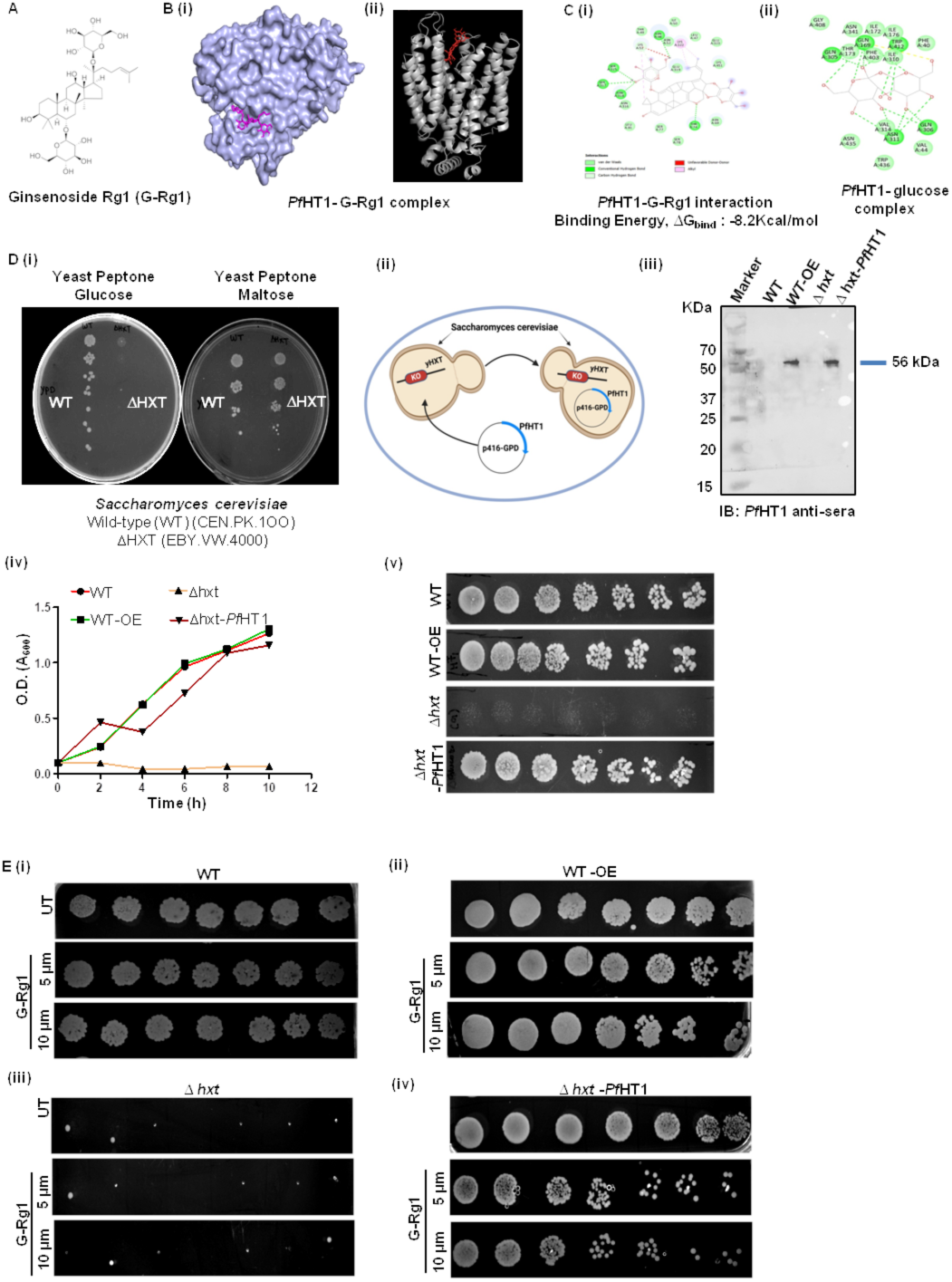
***In silico* interaction study of G-Rg1 with *Pf*HT1 and effect of G-Rg1 on growth of *Pf*HT1 complemented yeast. A.** 2D structure of G-Rg1 **B. (i)** *Possible architecture of the PfHT1-G-Rg1 complex* **(ii)** 3D structure of *Pf*HT1 highlighting the relevant catalytically active site and binding of G-Rg1 **C (i)** *In silico* interaction analysis of G-Rg1 with *Pf*HT1. G-Rg1 interacts via H-bonds (n =5) with ΔG bind of -8.2 kCal/mol. G-Rg1 primarily engaged with a pocket formed by residues Asn48, Lys51, Thr74, Ser315, Ser317, Asn318, and Glu319 **(ii)** *In silico* interaction analysis of *Pf*HT1 in complex with di-glucose indicated that the glucose interacts with residues Ile172, Thr173, Ile176, Gln169, Gln305, Gln306, Ile310, Asn311, and Val314, which is different from the G-Rg1 binding site. H-bonds. Bond lengths (Å) are highlighted as dashed lines **D.** Complementation of *Pf*HT1 in yeast mutant strain. **(i)** Culture of WT (CEN.PK2-1C) and Δ*hxt* (EBY.VW.4000) on yeast peptone dextrose (YPD) and yeast peptone maltose (YPM) plates **(ii)** Schematic representation of *Pf*HT1 complementation strategy in yeast **(iii)** Immunoblot showing the expression of *Pf*HT1 in WT and mutant yeast strains transformed with *Pf*HT1-p416-GPD. Expression was not observed in WT and mutant transformed with only p416GPD. Cultures of WT, Δ*hxt*, and their *Pf*HT1 complemented counterparts were harvested and adjusted to an optical density, A_600_ of 0.1 using sterile YPD media and incubated at 30°C. A_600_ of same cultures was spectrophotometrically measured after every 2 h to quantitate the growth of WT and complemented yeast cells. Subsequently, serial 10-fold dilutions of each culture at A_600_ of 0.1 were spotted onto YPD agar plates and incubated at 30°C for 2 days. Complementation with *Pf*HT1 rescues Δ*hxt* cell growth as depicted using growth curve **(iv)** and spot assay **(v)**, however no significant difference in growth of WT-*Pf*HT1 (OE) strain was observed. Upon complementing *Pf*HT1in Δ*hxt* strain, restoration of cell growth was observed. **E.** Effect of G-Rg1 on the growth pattern of Δ*hxt* strain complemented with *Pf*HT1. Spot-assay showing growth of yeast cells spotted following 2 days of incubation. Each culture was harvested and adjusted to A_600_ = 0.1 using sterile YPD medium. The particular solution was then serially diluted 10-times and each dilution suspension was then spot on solid YPD medium and incubated for two days at 30°C. Interestingly, *Pf*HT1-complemented yeast exhibited reduced growth in the presence of G-Rg1 as compared to the control (untreated). However, no effect of G-Rg1 on growth of WT yeast strain was reported, confirming the *Pf*HT1specificity of G-Rg1.

Interestingly, the binding pocket for G-Rg1 is distinct from the known di-glucose molecular binding pocket of *Pf*HT1. The known di-glucose binding site of *Pf*HT1 consists of interactions with residues Ile172, Thr173, Ile176, Gln169, Gln305, Gln306, Ile310, Asn311, and Val314 **[Figure 1C (ii)]**. This lack of overlap in binding sites suggests that G-Rg1 may not directly compete with glucose for binding to *Pf*HT1, but rather may exert its effects through an allosteric mechanism or by inducing conformational modifications in the protein. The strong binding energy of -8.2 kcal/mol for GRg1 coupled with its extensive hydrogen bonding network within the binding pocket, provides compelling evidence for its potential as a novel ligand for *Pf*HT1.

### *Pf*HT1 complementation in yeast mutant restores cell growth

For elucidating the functional expression of membrane transporter proteins including glucose transporters from diverse sources, yeast has proven to be an effective model system. In line with this, Boles and Hollenburg engineered the EBY.VW4000 yeast strain, a hexose transporter-deficient (*hxt^0^*) mutant, in which all hexose transporter-encoding genes as well as other transporters capable of hexose uptake have been deleted (19). EBY.VW4000 cannot grow on medium containing just glucose, fructose, or mannose, and grows poorly on galactose. For propagation, the *hxt^0^* strain is routinely cultured on maltose, a disaccharide transported via specific maltose symporters (encoded by the *MALx1* loci). Thus, the hxt^0^ strain provides a perfect platform to clone and characterize hexose transporters from other species, or to replace the function of indigenous transporters. Here in, to validate the functional role of *Pf*HT1 in *Pf* parasite*s*, we used an orthologous yeast model, *S. cerevisiae* [**Figure 1D (i) and (ii)**]. We effectively complemented the EBY.VW4000 strain with the *Plasmodium* hexose transporter, *Pf*HT1. The complementation was confirmed by immunoblotting using *Pf*HT1 anti-sera [**Figure 1D (iii)**]. When *Δhxt* yeast strain was complemented with *Pf*HT1-p416 GPD the growth of the complemented yeast re-established in glucose medium as compared to the *Δhxt* mutant strain, as shown by the growth curve and spot assays [**Figure 1D (iv) and (v)**]. CEN.PK2-1C was used as a wild-type (WT) control. The *Pf*HT1-p416 GPD complemented strain was used to identify the *Pf*HT1 inhibitory potential of G-Rg1.

### Ginsenoside Rg1 suppressed growth of complemented yeast by inhibiting *Pf*HT1

Having validated the role of *Pf*HT1 complementation in rescuing the growth of mutant yeast cells in the presence of glucose, the next objective was to determine whether G-Rg1 could specifically target *Pf*HT1 in the yeast complementation model. To address this, CEN.PK2-1C (WT), *hxt*Δ, and complemented yeast strains (WT-*Pf*HT1-p416 GPD (WT-OE), and *Δhxt*-*Pf*HT1-p416) were cultured in presence and absence of G-Rg1 for 12 hours to assess growth patterns. Notably, spot assay analysis revealed that *hxtΔ*-*Pf*HT1-p416 complemented yeast mutants exhibited reduced growth upon G-Rg1 treatment **[Figure 1E]**. Furthermore, the growth of WT-OE was also inhibited compared to the WT strain. This effect is likely attributable to the drug-targeting activity of G-Rg1 against *Pf*HT1. These findings strongly suggest that G-Rg1 selectively inhibits *Pf*HT1, leading to impaired growth, potentially due to restricted glucose uptake, thereby disrupting metabolic homeostasis.

### Expression of *Pf*HT1 in intra-erythrocytic drug sensitive and resistant parasites

The expression of *Pf*HT1 was demonstrated both at the RNA and protein levels. Stage specific transcript levels of *Pf*HT1 in *Pf*3D7 and *Pf*Kelch13^R539T^ were detected by qPCR utilizing cDNA obtained from tightly synchronized cultures of rings, trophozoites and schizont-stage infected erythrocytes of both the strains. *Pf*HT1 encoding transcripts were amplified using specific primers which revealed higher expression of *Pf*HT1 in the drug-resistant *Pf* parasites compared to sensitive counterparts, predicting a role of *Pf*HT1 in the survival of resistant parasites **[Figure 2A (i)]**. 18S served as a housekeeping control to ensure equal loading of RNA from each developmental stage for RT-qPCR analysis Immunoblot analysis was employed to evaluate the expression of native *Pf*HT1 in parasite lysates from the *Pf*3D7 and *Pf*Kelch13^R539T^ strains by using mice raised anti *Pf*HT1 antibodies. Lysates were produced from mix-stage population of parasites and distinct bands corresponding to *Pf*HT1 were found at the predicted molecular weights in both strains. A strong band of about 56 kDa was seen, which corresponds to the anticipated molecular weight of the full-length *Pf*HT1 protein in *Pf* **[Figure 2A (ii)]**.

**Figure 2:**
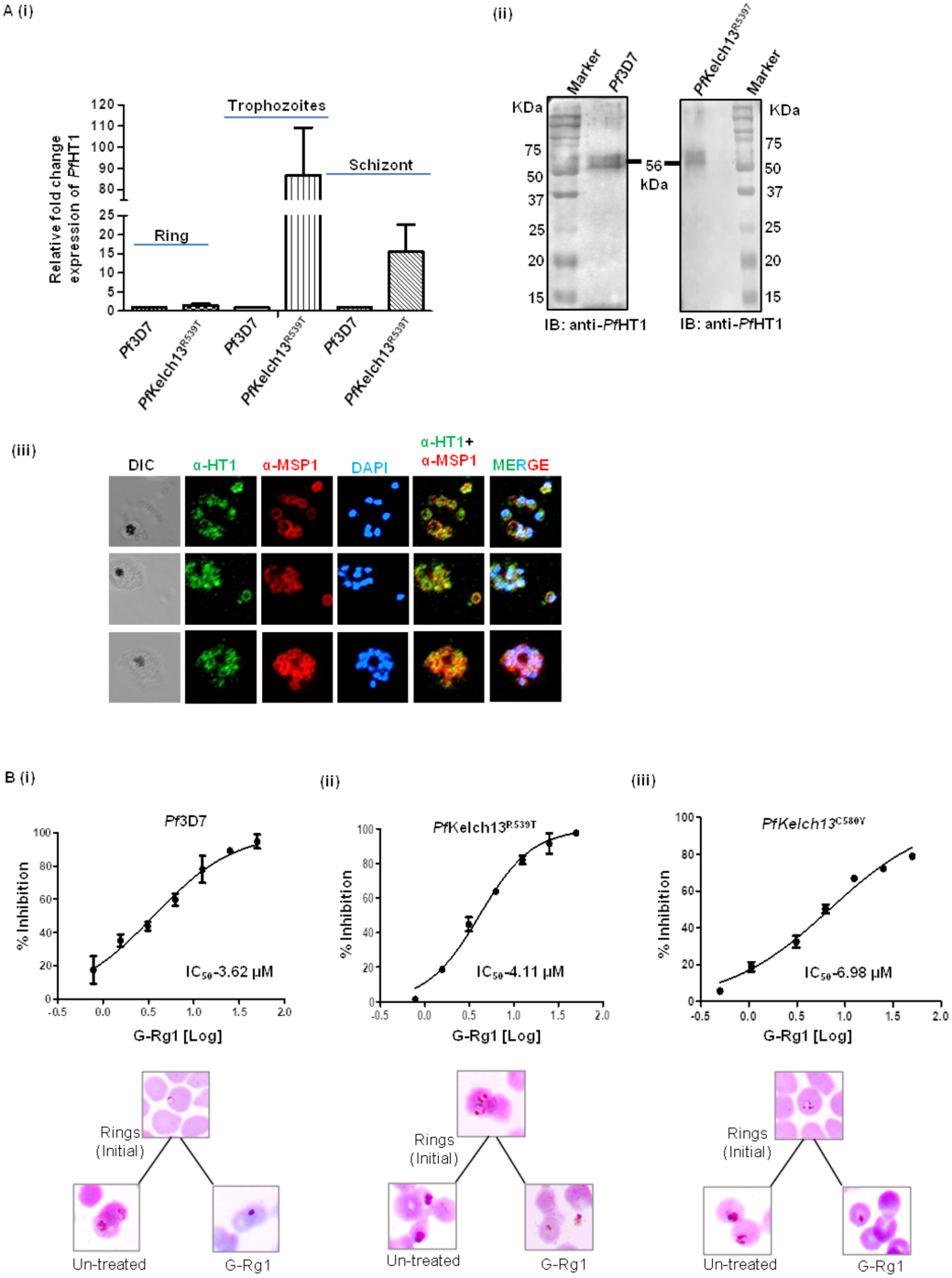
Expression and localization of *Pf*HT1 and effect of G-Rg1 on growth and progression of ART-sensitive and resistant strains of *P. falciparum* parasites. A (i) Quantitative RT-PCR analysis of *Pf*HT1 in cDNA derived from ring, trophozoites and schizonts stages of *Pf*3D7 and *Pf*Kelch13^R539T^. 18S rRNA transcripts were taken as a loading control **(ii)** Immunoblot using *Pf*HT1antisera showing *Pf*HT1 (̴̴̴̴ 56kDa) in mixed parasite lysates of *Pf*3D7 and *Pf*Kelch13^R539T^ **(iii)** Localization of *Pf*HT1 within mature schizont : green represents *Pf*HT1, red represents MSP-1. DIC represents Differential Interference Contrast. Parasite nuclei were stained with DAPI. IFA results depict that *Pf*HT1 is localized to the plasma membrane of parasite. **B.** Graphs representing concentration-dependent growth inhibitory effect of G-Rg1 on ART-sensitive *Pf*3D7 **(i)** and resistant *Pf*Kelch13^R539T^ **(ii)** and *Pf*Kelch13^C580Y^ **(iii)** strains, as evaluated by scoring Giemsa-stained parasites. The corresponding Giemsa-stained images of parasites pre- and post-66 h treatment at corresponding IC_50_ values of G-Rg1, showed the formation of pyknotic bodies (Scale bar = 2 μm).

### *Pf*HT1 localizes to the *Plasmodium* surface and colocalizes with MSP-1

To determine the cellular localization of *Pf*HT1 within the *Pf* parasite, fixed late stage parasite smears were immunostained with *Pf*HT1 antisera along with Merozoite Surface Protein-1 (MSP-1) antisera, a well-established marker of the merozoite surface. *Pf*HT1 was observed to be localized on the merozoite surface in punctated schizonts. Co-localization with MSP-1 (GPI-anchored membrane protein) indicates that *Pf*HT1 is localized at the parasite plasma membrane **[Figure 2A (iii)]**.

### Ginsenoside Rg1 suppressed the growth of intra-erythrocytic *P. falciparum* parasites

The *in vitro* treatment of ART-sensitive (*Pf*3D7) and resistant (*Pf*Kelch13^R539T^, *Pf*Kelch13^C580Y^) parasites with G-Rg1 demonstrated growth suppression with IC_50_ values of 3.62 μM, 4.11 μM and 6.83 μM, respectively [**Figure 2B (i), (ii) and (iii)**]. Furthermore, analysis of parasite development at 66 h post-treatment revealed that G-Rg1 impaired maturation into trophozoites in both drug-sensitive and resistant strains, instead inducing the formation of pyknotic bodies.

### Ginsenoside Rg1 impeded the growth of *P. falciparum* parasites maximally at the mature stages

During intra-erythrocytic development of *Pf*, parasite has to undergo a progression through several stages i.e., rings, trophozoite and schizonts. The growth-inhibitory capacity of G-Rg1 on various developmental stages was assessed using an *in vitro* stage-specific inhibition assessment. Parasites tightly synchronized at every stage were treated with G-Rg1 for 6 h subsequent to washing with iRPMI after every treatment period, resulting in parasites progressing in the absence of the drug for 66 h. A brief exposure (6 h) of parasites to G-Rg1 at different developmental stages demonstrated a stage-dependent chemo-suppressive effect **[Figure 3A**], with the highest inhibitory activity observed during the trophozoite and schizont stages (IC_50_ = 8.20 μM and 10.66 μM). IC_50_ value at the ring stage was found to be 17.95 μM. The increased efficacy of G-Rg1 at later stages can be linked with its highly active metabolic activity with increased glucose acquisition.

**Figure 3:**
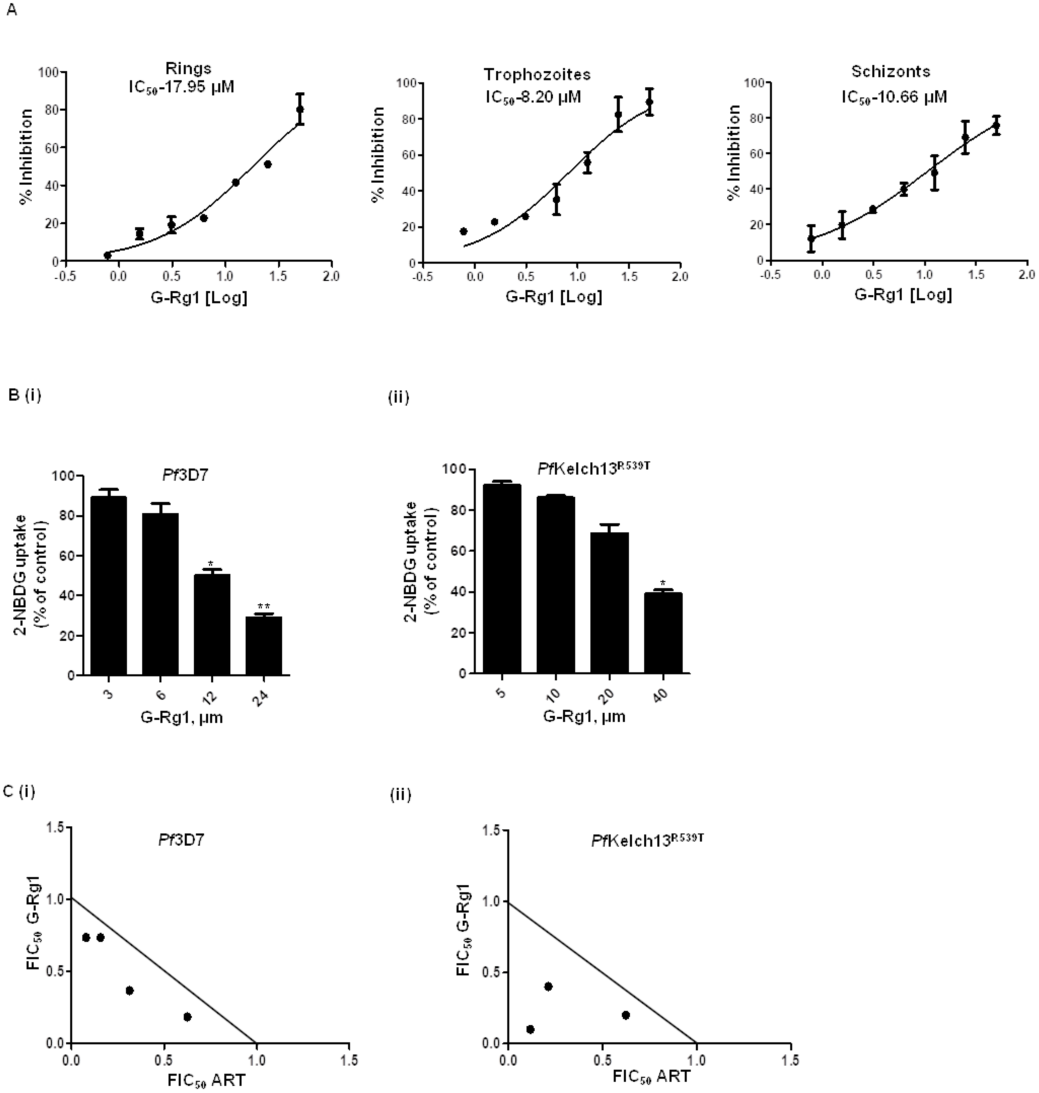
Stage specific growth inhibitory and glucose uptake blocking potential of G-Rg1 and its combinatorial effect with artemisinin. **A.** The graphs depicting the percent reduction in parasitemia by G-Rg1 at each parasitic development stage, along with the respective stage-specific IC_50_ values. The result depicts the mean of three independent experiments. **B.** G-Rg1 dose dependently inhibited the glucose uptake of both *Pf*3D7 **(i)** and *Pf*Kelch13^R539T^ **(ii)** parasites as assessed by 2-NBDG uptake assay. **C.** Combinatorial growth inhibitory effect of G-Rg1 and ART on *Pf*. Iso-bolograms depicting interaction between G-Rg1and ART against **(i)** ART-sensitive (*Pf*3D7) and **(ii)** resistant (*Pf*Kelch13^R539T^) strains of *Pf*. Mean FIC50 of ART and G-Rg1 were taken on the x-and y-axis, respectively. Results are representative of three independent experiments.

### Ginsenoside Rg1 reduced the glucose uptake of parasites

The effect of G-Rg1 on glucose acquisition of parasites was evaluated using a fluorescent probe, 2-NBDG. G-Rg1 dose dependently suppressed the glucose uptake of both drug sensitive and resistant parasites, thereby confirming its anti-malarial effect via inhibiting *Pf*HT1 [**Figure 3B (i) and (ii)**]. Depriving the parasites of their prime energy source can disrupt their metabolic signaling, resulting in cell death.

### Ginsenoside Rg1 and artesunate synergistically inhibited the growth of *Pf* parasites

Combination therapy is an effective strategy currently employed to tackle resistant malaria, involving the use of two or more therapeutic agents in conjunction. Checkerboard combination assay was used to determine the nature of *in vitro* interaction of G-Rg1 and ART on parasite growth. A synergistic effect was observed in the *Pf*3D7 strain at some concentrations, with an FIC₅₀ index of <1 **[Figure 3C (i)]**. Similarly, in ART-resistant parasites, G-Rg1 significantly enhanced ART efficacy, demonstrating a comparable synergistic interaction **[Figure 3C (ii)]**. Notably, G-Rg1 potentiated ART activity in both ART sensitive and resistant parasites, highlighting its potential as a combination partner for ART.

### Ginsenoside Rg1 enhanced antimalarial potency of dihydroartemisinin against resistant parasites

RSA is a reference assay to measure ART resistance against *Pf in vitro* setting. ART resistance is correlated with the reduced susceptibility of ring-stage parasites to DHA leading to delayed parasite clearance with ART monotherapy or 3-days ACT course in clinic. We evaluated the efficacy of G-Rg1 alone, and in combination with DHA on ART-resistant (*Pf*Kelch13^R539T^) parasites. Early ring-stage parasites treated with DHA 700 nM, various concentrations of G-Rg1 alone as well as in combination with DHA 700 nM exhibited normal progression of untreated parasites to mature trophozoites, whereas G-Rg1 and DHA-treated cultures displayed unhealthy and dead parasites [**Figure 4A(i)**]. Following incubation of 66 h, a significant reduction in parasite survival was observed in G-Rg1 treated parasites as compared to DHA-treated ones.

**Figure 4:**
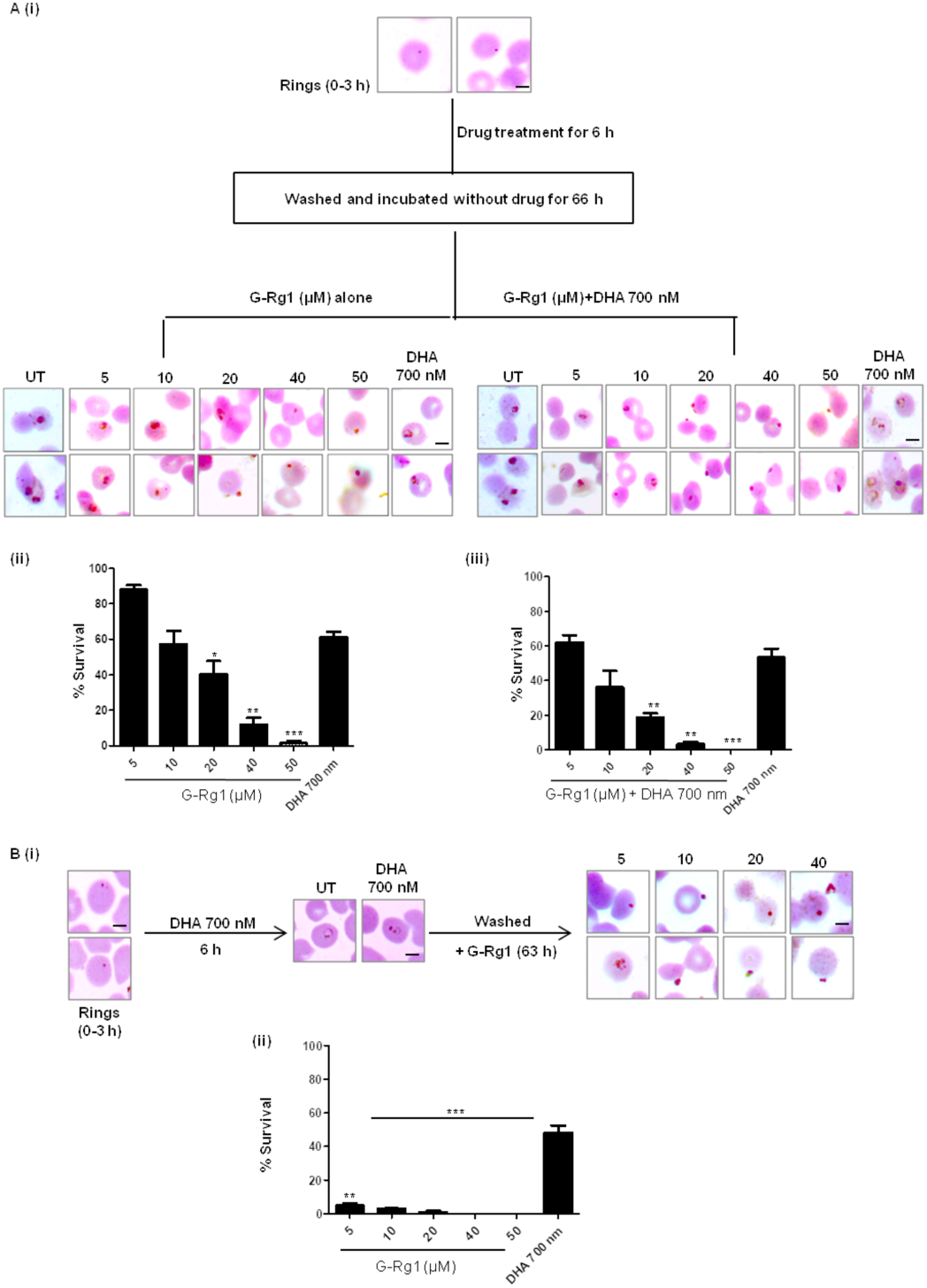
**G-Rg1 reduced the survival of DHA-resistant ring stage resistant parasites alone and in combination with DHA**. **A (i)** Schematic representation showing light micrographs of Giemsa-stained infected RBCs incubated with DHA and G-Rg1 alone as well as in combination with DHA. Bar graphs representing percentage survival following treatment with DHA and G-Rg1 alone **(ii)** and in combination **(iii)** The combination of G-Rg1 with DHA alleviates the antiparasitic activity of DHA against ring stages of ART-resistant parasites. **B (i)** Schematic diagram showing modified form of RSA, wherein ART-resistant ring staged parasites were treated with DHA followed by treatment with various concentrations of G-Rg1. The surviving parasites were scored and percentage survival was calculated for each condition. **(ii)** G-Rg1 potently decreases the survival of DHA pretreated resistant rings when compared to DHA alone. Results represent mean of three independent experiments. (*, P < 0.05, **, P < 0.01)

Notably, G-Rg1 treatment at 40 μM (8×IC₅₀) and 50 μM (10×IC₅₀) led to over 85% reduction in parasite survival, compared to a 48% reduction with DHA (700 nM) [**Figure 4A (ii)**]. Notably, G-Rg1 and DHA in combination increased the potency of DHA by lowering parasitemia by 70– 95% at different G-Rg1 concentrations [**Figure 4A (iii)**]. We further evaluated the inhibitory efficacy of G-Rg1 against resistant parasites that survived DHA treatment [**Figure 4B (i) & (ii)**]. The percentage of parasite survival (DHA-pretreated rings) reduced in the presence of G-Rg1, indicating that G-Rg1is powerful in clearing resistant parasites that survive post-DHA treatment, thus predicting it as a potential partner candidate for combination therapy.

### Ginsenoside Rg1 impeded parasite growth *in vivo*

*In vivo* antiplasmodial activity of G-Rg1 alone as well as in combination with ART was demonstrated using *P. berghei* ANKA-infected mice model **[Figure 5A (i)].** Infected mice were orally administered G-Rg1 (20 mg/kg), ART (60 mg/kg), G-Rg1 (10 mg/kg) + ART (30 mg/kg) and PBS as vehicle control for 5 consecutive days. The blood smears from tail vein were made regularly to estimate the parasitemia. G-Rg1 treated mice exhibited a 60% decrease in parasite burden compared to the vehicle control [**Figure 5A (ii) and (iii)**]. Additionally, mice treated with G-Rg1 exhibited a survival duration exceeding 15 days, in contrast to vehicle-treated controls, which succumbed within 10 days following infection [**Figure 5A (iv)**]. Notably, the combination of G-Rg1 and ART prominently suppressed parasite burden and extended the mean survival rate of infected mice. Mice receiving the combination therapy (at half doses of each drug) were alive even after 21 days. These preclinical findings highlight G-Rg1 as a potential partner drug for ART.

**Figure 5:**
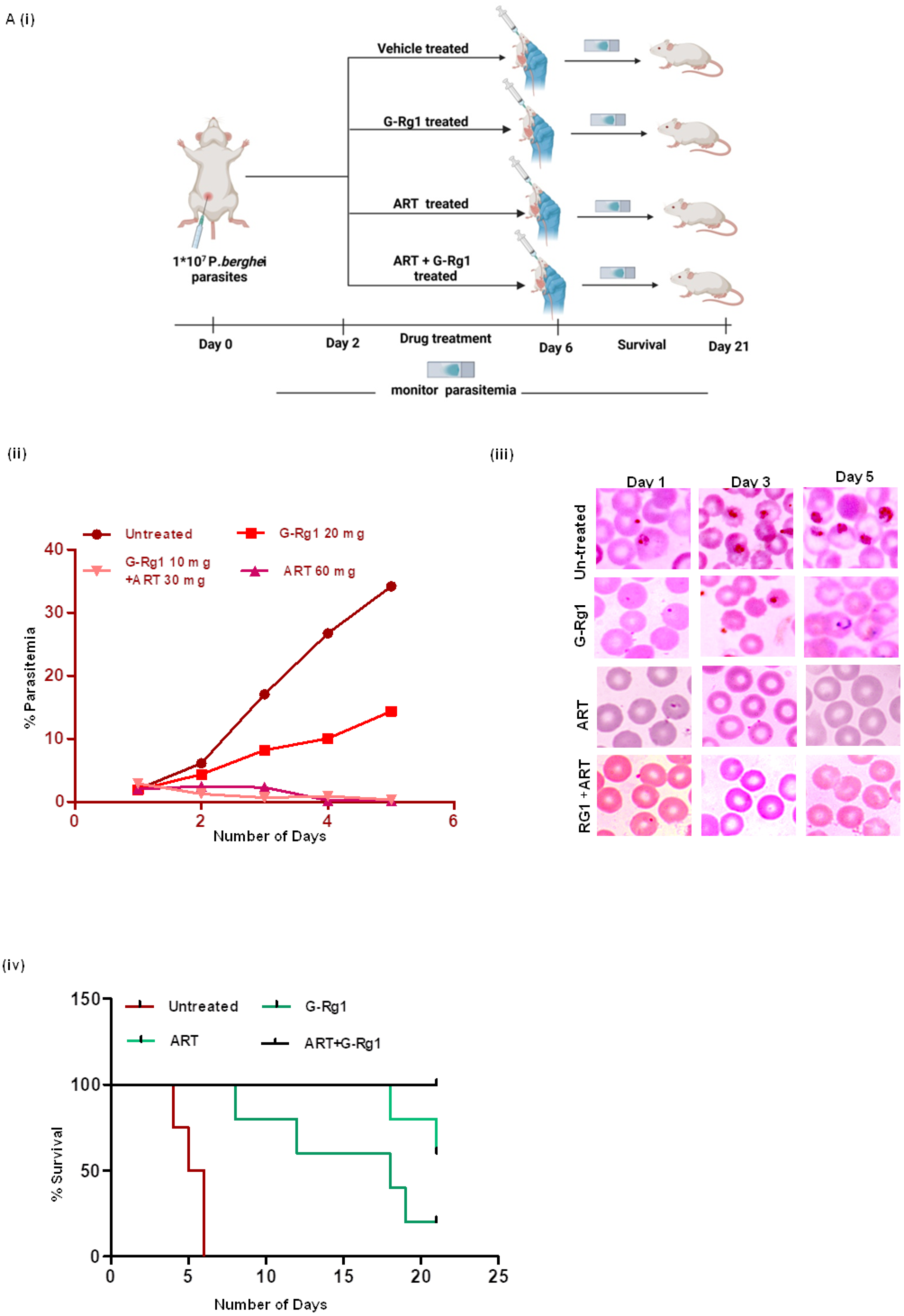
Effect of G-Rg1 on parasite growth in *in vivo* malaria model. **A** (i) Schematic representation of malaria induction and *in vivo* drug treatment regimen, vehicle-treated mice (n = 5) served as a control. **(ii)** Graph showing routine percent parasitemia of untreated, G-Rg1-treated, ART-treated and combination-treated infected mice. **(iii)** Micrographs of Giemsa-stained blood smears of control and treated mice, indicating parasite load in mice from each group**. (iv)** Plot showing the survival of untreated and drug-treated mice as observed for up to 21 days post-infection. G-Rg1, alone as well as in combination, significantly reduced the parasite burden and increased the survival of infected mice.

## DISCUSSION

The increasing prevalence of drug-resistant *Plasmodium* strains poses a significant challenge in malaria treatment (20), underscoring the need for investigation of new drug targets as well as innovative therapeutic strategies. *Pf*HT1 has emerged as a promising and highly selective therapeutic target to combat *Pf* infection (21, 22). As an essential membrane protein responsible for glucose uptake, *Pf*HT1 is the parasite’s sole facilitative glucose transporter throughout its intra-erythrocytic developmental stages (7). In contrast to the multiple redundant glucose transporters in human cells, *Pf* relies exclusively on *Pf*HT1, making it uniquely vulnerable to targeted inhibition (7, 9). Notably, phytochemicals represent a promising therapeutic leads for new antimalarial development. They have long served as a valuable source of bioactive compounds as most of the anit-malarials currently in use or historically significant trace their origins to natural products derived from plants (23). In line with this, our study identified a natural compound, G-Rg1, as a potential inhibitor of *Pf*HT1, through an *in silico* SBDD approach, and validated its *Pf*HT1 inhibitory potential using yeast complementation model. These complementation models offer a unique platform for functionally dissecting *Pf* membrane transporters and screening for selective inhibitors (24). Their simplicity, scalability, and specificity make them indispensable for preclinical antimalarial drug discovery pipelines, particularly when targeting essential nutrient or drug transporters like *Pf*HT1 (25, 26). Expression of *Pf*HT1 (codon-optimized) in *S. cerevisiae* mutant (EBY4000) yeast cells, that are null mutants for their endogenous hexose transporters (hxt) (19), restored their growth in glucose media, with respect to mutant (*Δhxt*) cells. Study results (growth curve and spot assay) revealed that the growth of the complemented line was completely dependent upon uptake of glucose through *Pf*HT1, confirming substrate specificity of complemented cells. *Pf*HT1 selectivity of G-Rg1 was also determined using the same complemented system. Interestingly, we observed that G-Rg1 decreased the growth of *Pf*HT1-complemented yeast cells providing evidence for the its specificity to *Pf*HT1.

Furthermore, the expression of *Pf*HT1 was validated at both the transcript and protein levels during the intra-erythrocytic stages of the *Pf* parasite, using quantitative RT-PCR and imunoblotting. Interestingly, stage specific mRNA expression results revealed an increased expression of *Pf*HT1 in resistant *Pf* parasites (*Pf*Kelch13^R539T^) compared to drug sensitive (*Pf*3D7) ones. This observation highlights a potentially underappreciated facet of resistance biology namely, the contribution of metabolic adaptations to drug tolerance and survival. The elevated expression of *Pf*HT1 observed in resistant parasites suggests a compensatory mechanism wherein parasites increase glucose uptake to sustain energy production under drug-induced stress. From a therapeutic perspective, these findings reinforce the rationale for targeting parasite metabolism in resistant *Plasmodium* parasites.

In line with this, our study results reported that G-Rg1 exhibited significant *in vitro* growth inhibitory effects on drug sensitive (*Pf*3D7) and resistant (*Pf*Kelch13^R539T^, *Pf*Kelch13^C580Y^) parasites, with an IC_50_ of 3.62, 4.11 and 6.83 μM, respectively. Notably, G-Rg1 had the capacity to decrease glucose absorption by parasites at the therapeutically active concentration, as determined by 2-NBDG uptake assay. Our findings align with the previous studies that emphasized that targeting glucose transport in parasites can starve them, leading to reduced growth and proliferation (12, 21).

The combination of *Pf*HT1 inhibitors with ART represents a rational and promising strategy to counteract ART resistance by exploiting the parasite’s metabolic vulnerabilities. As drug resistance continues to rise, such metabolism-targeted combinations may provide a vital tool for sustaining the efficacy of antimalarial therapies. Interestingly, we observed a synergistic anti-malarial association between G-Rg1 and DHA in both the strains, as determined by checkerboard combination assay.

Clinically, ART resistance is characterized by a delayed parasite clearance, specifically a half-life of ≥5 hours in peripheral blood following treatment. RSA is regarded as the gold standard assay for assessing ART resistance in *in vitro* settings (27). This assay requires tight synchronization of parasites to the 0–3 hour ring stage, a critical window that enables discrimination between ART-sensitive and ART-resistant strains (28). We employed RSA to evaluate the effect of G-Rg1 on the ART-resistant *Pf* parasites (*Pf*3D7^K13^R539T^). Our results demonstrated that G-Rg1, both as a monotherapy and in combination therapy with DHA, potently suppressed the growth of ART-resistant parasites. Furthermore, G-Rg1 significantly eliminated parasites that had survived prior DHA exposure. These findings further highlight the role of *Pf*HT1 in survival of ART resistant parasites.

Evaluating the *in vivo* efficacy of a compound is vital for the identification of promising lead candidates during the preclinical stage of drug development. We evaluated the *in vivo* anti- malarial activity of G-Rg1 in *P. berghei*-infected mice. G-Rg1 treatment alone and in combination with ART reduced the parasitic load as well as enhanced the survival of the mice. Altogether, our study not only advanced our understanding of *Pf* metabolism but also opened new avenues for the development of antimalarial therapies. Additionally, exploring the drug’s effects on other stages of the *Pf* lifecycle may reveal broader applications in malaria treatment. This study supports ongoing efforts to manage drug resistance and mitigate the global burden of this life-threatening disease.

## MATERIAL AND METHODS

### Screening of drug library for a potential inhibitor of *Pf*HT1

By utilizing an *in silico* therapeutic repositioning method, we screened the Drug Bank database of pharmacologically active compounds for similarities with the specific *Pf*HT1 inhibitor, C3361. Notably, C3361, a glucose analog, has been shown to moderately inhibit *Pf*HT1 and impede the survival and proliferation of intra-erythrocytic parasites *in vitro* (12). We retrieved Structural Data Format (SDF) files for C3361 and the Drug bank compounds from PubChem, which provides open access to chemical information and biological activity data. We performed structural superimposition of the drug bank library compounds with C3361 using Discovery Studio Visualizer v20.1.0.19295, generated by Dassault Systèms Biovia Corp., to evaluate structural similarity for each of the compounds with C3361 as a reference.

### *In silico* interaction analysis of Ginsenoside-Rg1 with *Pf*HT1

*In silico* molecular docking analysis was performed to investigate the binding interactions between the *Pf*HT1 and G-Rg1. SDF file of G-Rg1 was downloaded from PubChem and SDF format was changed to standard PDB format, subsequently followed by production of its energy-minimized 3D-structural structure using Chem3D Pro 12.0, as described earlier. The crystal structure of *Pf*HT1 (PDB ID: 6M20) was retrieved from the Protein Data Bank and prepared for docking using AutoDock Tools. AutoDock Vina was employed for the docking simulations due to its efficient search algorithm and accurate scoring function. A blind-docking approach was chosen to determine similarity in the docking sites of G-Rg1 and glucose. A grid box encompassing the entire *Pf*HT1 protein was defined. The dimensions and center of the grid box were adjusted to include all potentially interacting residues. Docking parameters were optimized to balance computational efficiency and thoroughness, with the exhaustiveness set to 8 and the number of output poses set to 9. Post-docking analysis was performed using Discovery Studio Visualizer. The best-scoring poses for each ligand was selected based on binding energy and visual inspection of the protein-ligand interactions. Interacting residues were identified through the molecular interactions in the binding pocket. Two-dimensional interaction diagrams were generated for G-Rg1 and glucose to visualize key protein-ligand contacts, including hydrogen bonds, hydrophobic interactions, and π-stacking.

### Generation of *Pf*HT1 complemented yeast strain

The Complementary Determining Sequence (CDS) that encodes *Pf*HT1 was codon-optimized for expression in yeast (GenScript Biotech, United States). *Pf*HT1 complementation in *Saccharomyces cerevisiae* (*S. cerevisiae*) yeast strains CEN.PK2-1C (Wild-type. WT) and EBY.VW.4000 (Hexose transporter mutant strain: Δhxt1-17) (29), was performed by transforming codon optimized *Pf*HT1-p416-GPD construct and p416-GDP (vector only) in these strains using Frozen-EZ Yeast Transformation IITM kit (Zymo Research) in accordance with manufacturer’s protocol, and plated on yeast nitrogen base (YNB) agar plate (supplemented with 2% glucose and/or maltose and 1×amino acid mix without uracil) as selective media, and cells were cultivated at 30°C. *S. cerevisiae* yeast strain CEN.PK2- 1C served as WT control. The complementation of *Pf*HT1-p416-GPD was further confirmed by immnuoblotting using *Pf*HT1 specific primary antibody.

### Growth analysis in *Pf*HT1 complemented yeast

To evaluate the growth dynamics of *Pf*HT1 complemented yeast mutant strains, we pre-cultured WT, WT-*Pf*HT1-p416 GPD (WT-OE), Δhxt and Δhxt-*Pf*HT1-p416 GPD strains in yeast peptone dextrose (YPD) media (1% yeast extract, 2% glucose and 2% peptone), at 30**°**C. Subsequently, the pre-culture was diluted, using YPD media, to an optimum optical density (A_600_) of 0.1 before being employed as the primary culture. Every 2 h, the A_600_ was quantified spectrophotometrically to determine the growth of WT and complemented yeast cells. Furthermore, after 24 h of incubation period, each culture was collected and corrected to A_600_ = 0.1 using YPD media. These solutions were then serially diluted ten times, and each diluted mixture was placed on YPD solid plates and cultured for two days at 30°C.

### Assessment of Ginsenoside-Rg1 selectivity for *Pf*HT1

To evaluate the specificity of G-Rg1 towards *Pf*HT1, WT, WT- *Pf*HT1-p416 GPD (WT-OE), Δhxt and Δhxt-*Pf*HT1-p416 strains were grown overnight in YPD media, until adequate growth was accomplished. Consequently, yeast cultures were diluted with sterile YPD medium to an optical density (A_600_) of 0.1 and maintained in the presence or absence of 5 and 10 μM G-Rg1 for 12 h at 30°C. After incubation, the cultivated cell suspension was serially diluted 10-fold and spotted onto YPD agar plates and cultured at 30°C for two days.

### Parasite culture

*Pf* laboratory-adapted strains *Pf*3D7, *Pf*Kelch13^R539T^ and *Pf*Kelch^C580Y^ were cultured *in vitro* in accordance with the standard protocols of Trager and Jansen (30). Briefly, parasites were cultured O+ erythrocytes (Rotary Blood Bank, New Delhi) at 2% hematocrit in RPMI 1640 medium (Gibco, USA), supplemented with Sodium bicarbonate (2 g/l, Sigma Aldrich, USA), Hypoxanthine (50 mgL^-1^, Sigma Aldrich, USA), Albumax I (0.5%, Invitrogen, USA), and Gentamicin (10 mg/L, Sigma Aldrich, USA), at 37°C in a mix gas setting (90% N_2,_ 5% CO_2_ and 5% O_2_). Parasite culture was regularly monitored by preparing thin smears, fixing them with methanol (Merck) and staining with Giemsa solution (5%, Sigma Aldrich, USA). Giemsa- stained parasites were visualized using a compound microscope (Olympus, Tokyo) with oil immersion at 100X magnification. Synchronization of parasites at the ring stage was obtained by treating them with sorbitol (5%, Sigma Aldrich) in two successive cycles, following established protocols.

### Stage specific analysis of *Pf*HT1 using RT-PCR

Stage-specific expression of *Pf*HT1 was confirmed through reverse transcription qPCR. The cDNA from synchronous cultures of *Pf*3D7 and *Pf*Kelch13^R539T^ strains at the ring, trophozoite, and schizont stages was used to amplify transcripts corresponding to *PfHT1* and *18S* rRNA genes using the following set of primers, *Pf*HT1: Forward: 5’- GGTGCTGTGTTAGGATGTGGT -3’; and, Reverse 5’- ACACCATACGCACCCTTCTT -3’, and *Pf*18S: Forward: 5’-CCGCCCGTCGCTCCTACCG-3’; and, Reverse 5’- CCTTGTTACGACTTCTCCTTCC-3’. The resulting data was analyzed using Step-One software (Applied Biosystems) by calculating the comparative Ct values for the reactions.

### Detection of *in vivo* expression of *Pf*HT1 in parasites

To evaluate *Pf*HT1 expression in *Pf* parasites, late-stage *Pf*3D7 and *Pf*Kelch13^R539T^ parasites were extracted using saponin (5%, Sigma Aldrich, USA).The resulting parasite pellet was lysed with RIPA buffer supplemented with protease inhibitor. The lysate was then resolved using SDS-PAGE (12%), followed by transfer onto a nitrocellulose membrane. The blot was probed with *Pf*HT1 anti-sera diluted to 1:500 for 1 h at room temperature (RT), followed by probing with an HRP-conjugated anti-mice secondary antibody (1:5,000; Sigma-Aldrich) at RT for 1 h.

Immuno-fluorescence Assay (IFA) was conducted on *Pf*3D7 parasites to detect the localization of *Pf*HT1 in *Pf* parasite, as described earlier (31). The mature-stage parasite smears were probed with anti-*Pf*HT1 (1:200) and anti-MSP-1 antibody (1:300), followed by incubation with secondary anti- bodies (anti-mice Alexa fluor 488 and anti-rabbit AF 546). The slides were treated with DAPI antifade (Invitrogen) and pictographed using confocal microscope (NIKON Corporation).

### Growth inhibition assay

To determine the anti-malarial potential of G-Rg1 and to assess its half-maximal drug inhibitory concentration values (IC_50_) against *Pf*3D7, *Pf*Kelch13^R539T^, *Pf*Kelch13^C580Y^ asexual blood stage parasites, synchronized *Pf* infected ring stage erythrocyte cultures at a parasitemia of 0.8% and a hematocrit of 2% were incubated with different concentrations [50-0.58 μM] of G-Rg1 for 66 h. Untreated and positive (artesunate) controls were cultivated and processed under identical conditions. To assess the inhibitory effect of G- Rg1 on parasite growth, parasitemia was estimated by manually scoring the number of infected cells in an approximated 2000 erythrocytes, and % inhibition was computed relative to the untreated control employing this formula.

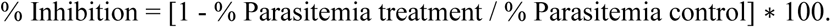

The IC_50_ values of G-Rg1 in *Pf*3D7, *Pf*Kelch13^R539T^ and *Pf*Kelch13^C580Y^ were estimated by plotting percent inhibition values versus log concentration of compound using Graph-pad PRISM software.

### Stage specific growth inhibition assay

The stage-specific growth inhibitory effect of G-Rg1 on *Pf* parasites was evaluated following a standardized protocol (32). Briefly, synchronized parasites (*Pf*3D7) at 0.8% parasitemia were subjected to different concentrations of G-Rg1 (for 6h) at ring stage (6 to 12 h), trophozoite stage (28 to 34 h), and schizont stage (42 to 48 h). After the 6-hour incubation, G-Rg1 was eliminated by washing the culture with complete medium, and the parasites were cultured further till the trophozoite stage of next cycle. Untreated parasites were treated as control. The percentage of chemo-suppression was determined by comparing the parasitemia levels of drug-treated parasites with those of the untreated control

### 2-NBDG uptake assay

Glucose uptake by *Pf* parasites was quantified using 2-(N-(7-nitrobenz- 2-oxa-1,3-diazol-4-yl)amino)-2-deoxyglucose (2-NBDG), a fluorescent D-glucose analog (Sigma-Aldrich, USA), measured spectrophotometrically (33). Synchronized early trophozoite stage parasites were treated with G-Rg1 (IC_50,_ 2x IC_50_, 4x IC_50_ and 8x IC_50_) and kept for 6 h at 37 °C. The culture media was exchanged with RPMI (without glucose) supplemented with 2-NBDG (0.3 mM), accompanied by incubation at 37 °C for 30 min to facilitate the absorption of the glucose analogue. To end the 2-NBDG absorption process, the incubation mixture was removed and the cells were washed with RPMI media twice, followed by washing with phosphate buffered saline (PBS). Further, samples were employed to determine the intensity of 2-NBDG fluorescent parasites by spectrophotometer (Thermoscientific, USA). Data was plotted as percent decrease in glucose uptake with respect to control.

### Drug combination assay

The combinatorial effect of G-Rg1 and ART on *Pf* parasites was demonstrated in accordance with the checkerboard assay (34). In this experiment, two drugs are evaluated in double serial dilutions, and the effect of each drug is determined both as monotherapy and in combination. The interaction between the two drugs is then demonstrated either geometrically or algebraically. In brief, ring staged *Pf* parasites (*Pf*3D7 and *Pf*Kelch13^R539T^) at 0.8-1% parasitemia were exposed to varying concentrations of G-Rg1 (0.58– 20 µM) and ART (0.58–20 nM), both individually and in combination, for 66 hours following the checkerboard assay protocol. Untreated parasites were marked as control. Fractional inhibition concentrations (FICs) were calculated as reported earlier, with FIC values <1 indicating a synergistic interaction between the two compounds.

### Ring Survival Assay

To determine the efficacy of G-Rg1 on *Pf*Kelch13^R539T^ parasites, RSA was carried out as described previously (35). In brief, schizonts (percoll-purified) were cultured until the early ring stage (0-3 h post-invasion), and subjected to varying concentrations of G-Rg1, DHA (700 nM), and their combination for 6 h. After incubation parasites were washed and maintained for an additional 66 h. Parasitemia was determined by counting approximately 5000 erythrocytes on Giemsa-stained smears. Additionally, to evaluate the effect of G-Rg1 on survival of DHA-resistant parasites, early ring-stage (0–3 h) parasites were first treated with DHA (700 nM) for 6 h, followed by washing and subsequent exposure to G-Rg1 for 66 hours. Parasite survival was assessed by scoring Giemsa-stained smears.

### In vivo assay

The *in vivo* anti-malarial potential of G-Rg1, both as a monotherapy and in combination with ART, was evaluated in *P. berghei* ANKA-infected BALB/c mice (36). All experimental procedures were conducted in accordance with the guidelines established by the Institutional Animal Ethics Committee (IAEC) of Jawaharlal Nehru University (JNU, IAEC no. 38/2024). Mice obtained from Central Laboratory Animal Resources (CLAR), JNU were infected with 1x10^7^ parasites intra-peritoneally at Day 0 and segregated randomly into four groups. (n=5). On day 2, mice in different groups were administered G-Rg1 (20 mg/kg in PBS, p.o.), ART (60 mg/kg in PBS, p.o.), G-Rg1 (10 mg/kg in PBS, p.o.) + ART (30 mg/kg in PBS, p.o.) and PBS (vehicle control) respectively for 5 days (Figure. 2.1). Parasitemia was quantified each day from Giemsa-stained tail blood smears. The mean survival time (in days) for mice in each experimental group post-inoculation was calculated over a 21-day observation period.

### Statistical analysis

Statistical analyses were done using GraphPad 5 (GraphPad Software Inc.) and p-values were estimated by two-tailed Student’s t-test wherever required.

## ACKNOWLEDGMENT

S Singh gratefully acknowledges the financial support in the form of Indian Council of Medical Research (ICMR; NER/84/2022-ECD-I) and Department of Biotechnology (DBT), Government of India (Project No. IC-12044(11)/10/2021-ICD-DBT). S Shafi acknowledges Senior Research Fellowship from ICMR (45/48/2019-PHA/BMS). We highly acknowledge Prof. Eckhard Boles, University of Frankfurt for providing us with the yeast mutant stains.

## AUTHOR CONTRIBUTION

**Sadat Shafi:** Methodolgy, Validation, Formal analysis, Investigation, Data curation, Software, Writing-Original Draft, and Writing- Review and Editing. **Preeti Maurya:** Methodology and Investigation. **Ruby Bansal:** Methodology and investigation. **Jyoti Sharma**: Methodology and investigation. **Sai Kumar Mishra:** Methodology and Investigation. **Abul Kalam Najmi:** Methodology, Validation, Writing-Review and Editing. **Shailja Singh:** Conceptualization, Methodology, Validation, Formal analysis, Writing – Original draft, Writing- Review and Editing, Resources, Data curation, Supervision, Project administration, Funding acquisition

## DECLARATION OF COMPETENT INTEREST

The authors declare that they have no known competing financial interests or personal relationships that could have appeared to influence the work reported in this paper.

## DATA AVAILABILITY

Data will be made available on request.

